# Data-driven modeling of *E. coli* transcriptional regulation

**DOI:** 10.1101/2024.05.30.596718

**Authors:** Christopher G. Dalldorf, Griffith Hughes, Gaoyuan Li, Bernhard O. Palsson, Daniel C. Zielinski

## Abstract

The growth of bacterial gene expression datasets has offered unprecedented coverage of achievable transcriptomes, reflecting diverse activity states of the transcription regulatory network. Machine learning methods like Independent Component Analysis (ICA) can decompose gene expression datasets into regulatory modules and condition-specific regulator activities. Here, we present a workflow to utilize inferred regulator activities to construct quantitative models of promoter regulation in *E. coli*. Resulting models are validated by predicting condition-specific TF effector concentrations and binding site motif strength based on differential gene expression data alone. We show how reconstructed promoter models can capture multi-scale regulation and disentangle regulator interactions, including resolving the apparent paradox where *argR* expression is positively correlated with its regulon despite being a repressor. We applied the workflow for all regulator-linked components extracted by ICA, demonstrating the scalability of the workflow to capture the *E. coli* TRN. This work suggests a path toward systematic, quantitative reconstruction of transcription regulatory networks driven by the large-scale databases that are now available for many organisms.

## Introduction

The transcription regulatory network of bacteria is a critical determinant of cell state and responsiveness^1^, and largely involves trans-acting regulatory proteins which bind to promoter regions to promote or deter recruitment of RNAP. Binding sites and activities of key regulators such as transcription and sigma factors have been painstakingly determined over the years through experiments such as ChIP and gSelex^2^. Meanwhile, observations of transcription regulator states, in the form of gene expression datasets, have become increasingly available for diverse conditions. In *E. coli*, the scale of both TF binding identification and RNA-seq have been rapidly increasing, with only 121 of the estimated 300 TFs having any known gene targets in 2003^3^ compared to 232 TFs with gene targets with strong or confirmed confidence today^2^. The development of sequence analysis tools has enabled binding site identification purely through computational means^2^, including some tools which can additionally predict binding strength^4^. As of 2024, NCBI GEO contains 23,890 individual samples of RNA-seq for *E. coli*, while at the end of 2010 there were only 4,127 samples^5^. This abundance of data suggests that there exists the potential for an integrative understanding of transcription regulatory network function in *E. coli*.

TRN network inference from gene expression datasets is a classic problem in systems biology^6^. Approaches to infer transcriptional regulation from large datasets generally focus on identifying regulatory interactions and strengths rather than explicitly modeling the biochemical nature of the resulting regulation. These methods for TRN estimation contrast with another paradigm for network assembly, network reconstruction, which focuses on the bottom-up structured assembly of biological knowledge through manual curation^7^. Although transformative in modeling metabolism, the reconstruction workflow when applied to transcription regulatory networks to date has relied on experimental mappings of transcription units^9^ and computationally determined motifs^10^ and resulted in largely qualitative (Boolean)^8^ models. However, a key distinction of reconstructions compared to other knowledge bases such as encyclopedias^11^ is that they are structured to enable the generation of quantitative models whose predictive performance can be evaluated. Thus, there is the potential for data-driven and reconstruction based approaches to be unified within a quantitative framework to more accurately capture transcription regulation.

Independent component analysis (ICA) has been a successful approach for *de novo* inference of transcriptional modules from gene expression databases^12^. The resulting activities of these components offer an estimate of the regulator activity on a given condition. Presumably, with a knowledge of transcription factor regulons and sufficient measurements of the outcomes of regulation, one can create a model to directly connect expression to biophysical measurements of TFs such as concentration and **K_d_** values. This approach seeks to learn the function of TRNs from observing their behavior across conditions using methods like ICA which capture real regulon structure and activities, thereby mapping phenomenological parameters to mechanistic ones.

In this study, we develop a workflow to quantitatively reconstruct the TRN of *E. coli* by building mechanistic promoter models and parameterizing them utilizing large-scale gene expression data. We first define condition-specific regulon activities using ICA. We utilize known and inferred regulons to generate mechanistic models for all regulated promoters characterized to date. We then parameterize these models in a stepwise process, going from phenomenological ICA activities to fundamental biochemical parameters. We validate our approach by predicting effector metabolite concentrations and TF dissociation constants. The resulting models are used to disentangle complex regulation scenarios and interpret mutations. We extend this workflow to all regulons for *E. coli* with activities that can be inferred from ICA, representing a substantial fraction of all known regulation. This work lays the groundwork for a quantitative understanding of transcriptional regulatory networks in bacteria at a new scale.

## Results

### Constructing promoter models with data-inferred regulator activities

First, we describe the inference of condition-specific regulator activities from expression datasets. Various machine learning methods have been used to separate gene expression data into groups of genes forming co-regulated modules and regulator activities. Independent component analysis (ICA) has been demonstrated as one of the most effective methods based on the ability to capture experimentally-determined regulons. Independent components from gene expression analysis have been termed iModulons, or independently modulated groups of genes. The process of generating iModulons using ICA can be best understood as a blind-source separation of regulatory signals. ICA separates gene expression data (X) into groups of genes called iModulons (M) and the activity of these iModulons across the input samples (A) (**Figure 1A**). The latest version of iModulons produced for *E. coli* is the PRECISE1K dataset^13^ which includes over 1000 individual expression profiles and generates 201 iModulons that altogether account for 86% of known regulatory interaction.

**Figure 1.**
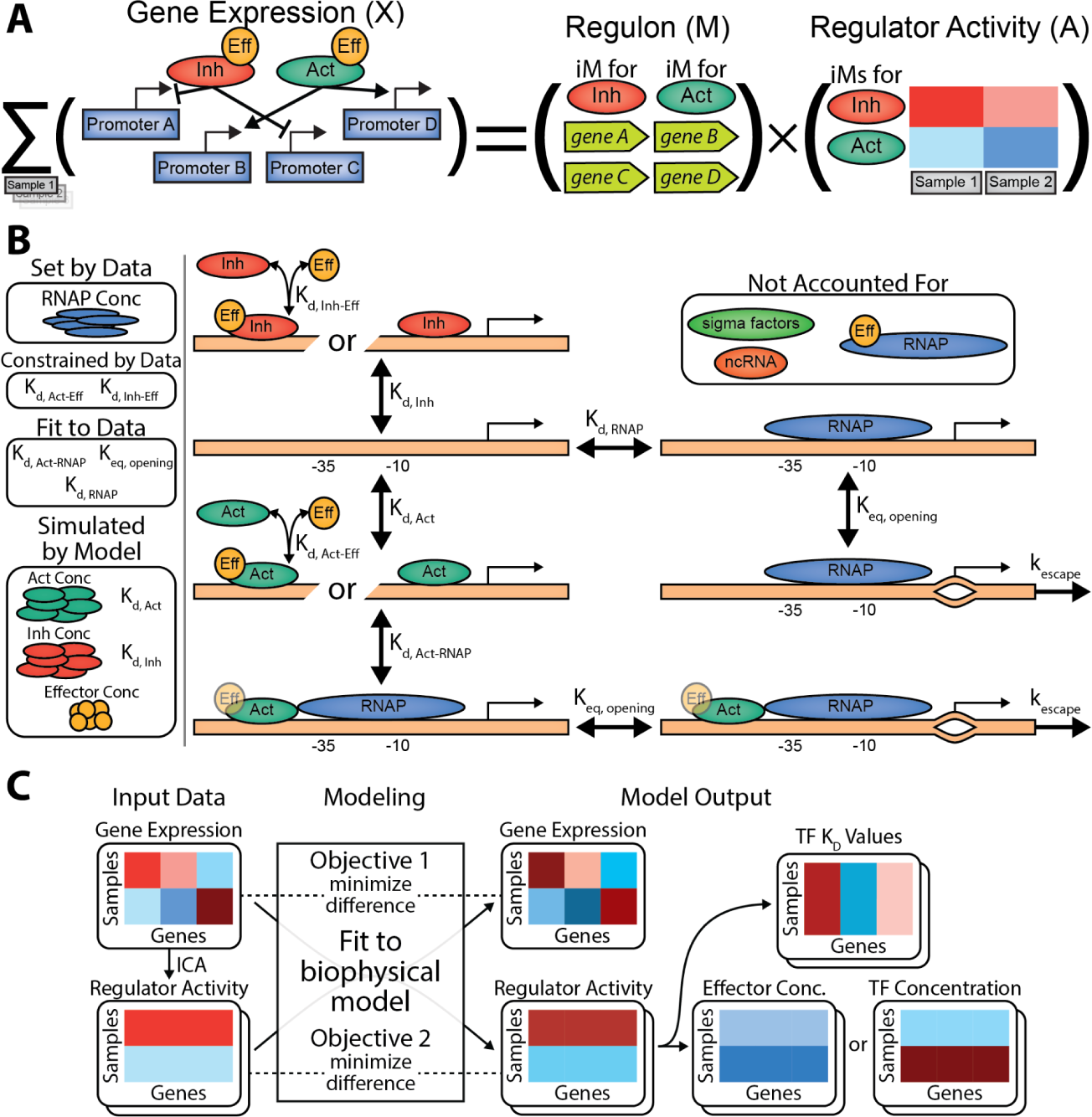
Workflow for data-driven construction of transcriptional regulatory models. **(A)** Application of independent component analysis to expression data in order to approximate regulatory activity. **(B)** Mechanistic promoter model along with biological constants set by the data, fit to the data, and simulated by the model. The different promoter states this study’s model incorporates are shown. Some relevant factors to gene expression are not modeled, such as sigma factors, non-coding RNA, direct RNAP binding effectors, and the concentration of genes. **(C)** Overall simplified workflow diagram for our model.

The premise of the modeling workflow is to use these iModulons as indicators of condition-specific regulatory activity, then subsequently infer a biophysical promoter model that matches both these regulator activities and resulting gene expression. The generation of this model involves inferring biological variables effector concentration, regulator concentration, and transcription **K_d_** values, providing a basis for independent validation of the generated models. RegulonDB, a compendium of regulatory binding sites, provides a knowledge base of what genes are regulated by what transcription factors^2^. This information, combined with inferred regulator activity of said transcription factors from ICA enables us to create a biophysical model of transcriptional regulation. To do this, we created a thermodynamic model of regulated transcription with different promoter states represented along with binding and rate constants (**Figure 1B**). We also model effector concentrations along with their binding constants to transcription factors which require effectors.

The mathematical equations connecting these promoter states to each other and to the expression values are outlined in **Methods: Mathematical Model Equations**. In summary, they are able to calculate expression values for a specific gene based on cActivator and cInhibitor which, respectively, refer to the regulatory activity of a promoter and a repressor. The equations also require several biological constants to be set, such as **K_d, RNAP_**, **K_eq, opening_,** and **K_d, Act-RNAP_**. These are set by calculating various possible solution sets that satisfy the underlying mathematical equations and picking the set that creates the best range of cActivator and cInhibitor values (see **Methods: Selection of Biological Constants**). Other values are hard set by the data itself, such as RNAP concentration, **K_d, Act-Eff_** and **K_d, Inh-Eff_** (see **Methods: Selection of Biological Constants**). Additional parameterization steps are further described in the methods.

Figure 1C gives a summary of the resulting workflow. With the mathematical model set and biological constants ready, we now need to generate the input cActivator and cInhibitor values. If a gene has only an inhibitor or activator, these can be directly solved as the number of equations and unknowns is equal. The ICA reconstructed expression value, M × A of the gene and the activating or inhibiting iModulon, is used as the expression value in the equation. If there is an inhibitor and activator for a gene, these are created using a genetic algorithm which creates a set of valid solutions to the expression equations and selects the solutions which best correlate with the activating and inhibiting iModulons. An additional greedy algorithm further optimizes these solution sets for input into the modeling software, GAMS (see **Methods: Genetic and Greedy Algorithm Optimization**). GAMS also requires the different activation types of the transcription factors involved, which can be a TF that is active without an effector, a TF that is active with a single effector molecule, or a TF that requires two copies of the same effector molecule to be active.

The process above is repeated for each gene contained in a specific activator-inhibitor pair we call a regulatory case. For example, all genes that are repressed by the Arginine iModulon and have no activating iModulon are grouped together as a regulatory case, which GAMS runs on independent of all other genes. The GAMS model optimizes for two criteria: 1) matching the input and output cActivator and cInhibitor values; and 2) matching the actual and predicted mRNA values. Additional details about the GAMS model are available in **Methods: GAMS Model**. The GAMS model outputs predicted mRNA values, GAMS optimized cActivator/cInhibitor values, and predicted biological constants for the regulators and promoter sites. Depending on the activation type of the associated transcription factors, these regulator-associated biological constants can include metabolite concentration, **K_d, Act_**, **K_d, Inh_**, and TF concentrations.

### Multi-scale model resolves counter-intuitive regulation of arginine biosynthesis genes

We first examined the ability of the established transcriptional regulatory modeling framework to capture the regulation of the ArgR regulon, which regulates the arginine iModulon consisting of arginine biosynthesis genes. We observed that expression of *argA*, along with the other genes repressed by ArgR, is positively correlated with *argR* expression in the PRECISE1k database (Figure 2A). This is counter-intuitive, as higher repressor expression logically should infer higher repressor activity and thus less expression. We also noticed that the Arginine iModulon has a much higher correlation to *argA* than *argR*. Thus, we hypothesized that the Arginine iModulon activity may is a better approximate for effective ArgR activity than *argR* expression, and may actually have an inverse relationship with effective ArgR activity due to the effect of the ArgR activator arginine, which was an unknown in the model. Figure 2C shows a summarized view of *argR* and *argA* expression, highlighting the important role that arginine concentration itself plays in said regulatory network.

**Figure 2.**
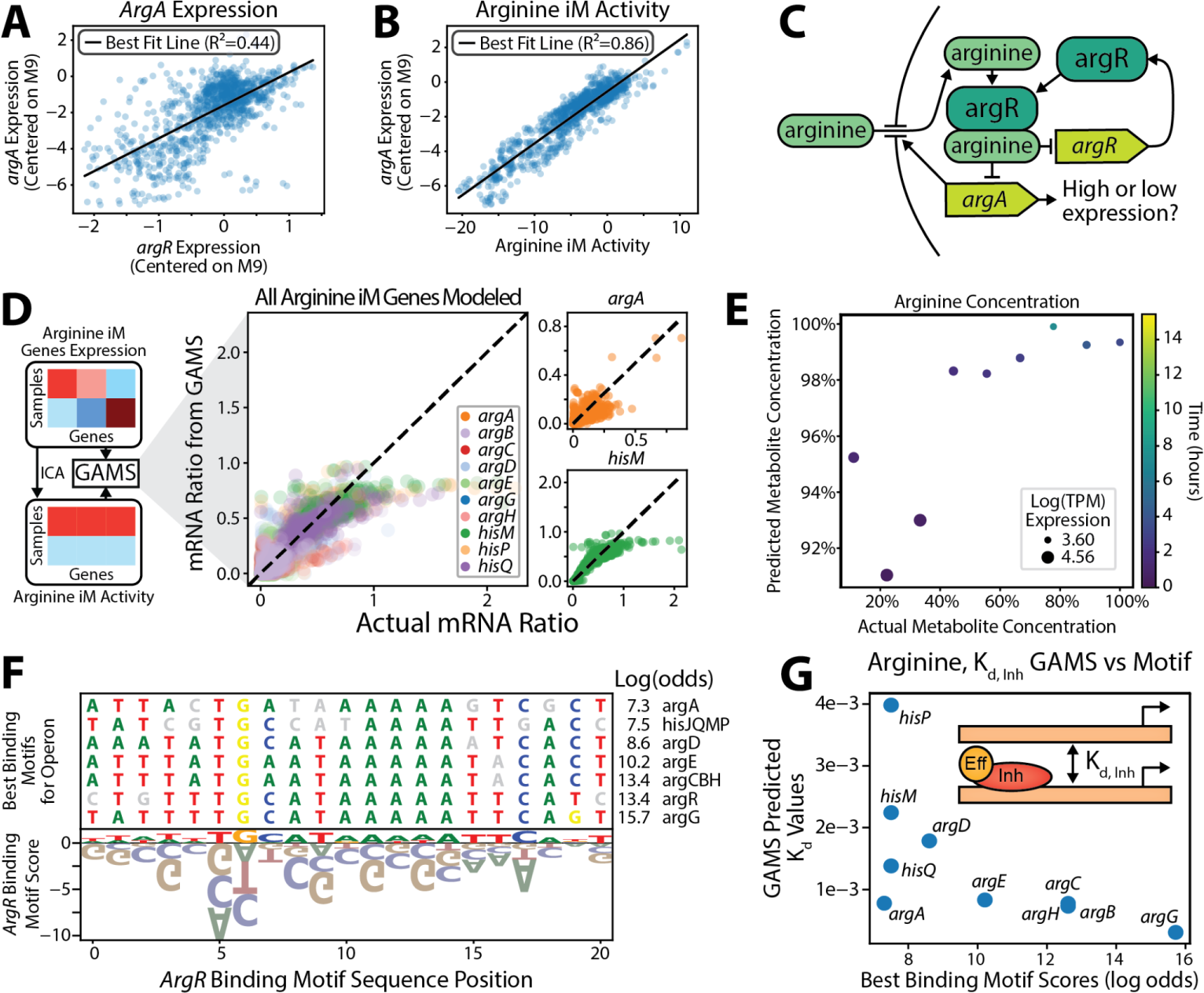
Transcriptional regulatory model captures multi-scale argR regulation. **(A)** *ArgA* expression is positively correlated with its repressor, *argR*. Samples are from PRECISE1K. **(B)** *ArgA* expression is highly positively correlated with the Arginine iModulon activity levels, showing it is a better indicator for regulatory activity than *argR*’s expression alone. **(C)** The regulatory network surrounding *argA* and *argR*, which is dependent on arginine concentration and also regulates various arginine processes including its import. **(D)** Inputs and outputs of GAMS model. GAMS is able to accurately predict gene expression. **(E)** Predicted and actual metabolite concentrations are highly correlated, which provides an external validation of the model. **(F)** *ArgR* binding motif alongside the most likely binding sites for the modeled operons. **(G)** Predicted GAMS **K_d_** values for *argR* binding are highly correlated with the sequence-based binding calculations.

Accounting for both ArgR and arginine levels, our model is able to use this inferred regulatory activity from the iModulon to accurately recreate gene expression values (Figure 2D). This workflow is able to untangle this complex and non-intuitive regulatory network by incorporating arginine concentration. The predicted arginine concentrations highly correlates with experimentally measured arginine values^14^ for a subset of conditions where these measurements were available (Figure 2E). This offers external validation of our workflow and highlights its ability to predict the metabolome using expression data.

In addition to validation by experimental data, we can also compare sequence-based predictions of *argR* binding strength for the modeled genes to their predicted **K_d_** values. Higher binding strength would mean less of the active transcription factor would be necessary for expression, thus meaning binding strength and **K_d_** values should be anti-correlated. We calculated the binding motif strength between *argR* and the promoter sites of the modeled genes (Figure 2F). This resulted in an expected highly negative correlation between predicted **K_d_** values and sequence-based predictions of binding strength (Figure 2G).

### Validation of models through prediction of effector concentrations

We next examined the PurR regulon, which regulates the Purine iModulon and has two different metabolite effectors, guanine and hypoxanthine^15^. We generated a model that accounts for both effectors interchangeably, due to unknown binding constants, and utilizes total purine as a variable (Figure 3A). For data from a joint expression and metabolite experiment^14^, the resulting model is able to accurately predict this summed concentration at early time-points post glucose starvation. However, the model fails to predict purine concentrations during late starvation (Figure 3B). This failure could be due to a handful of factors, including the fact that the model does not yet account for sigma factors or direct RNAP effectors, as this data is primarily from stationary phase samples where *rpoS* and ppGpp play large roles in gene regulation^16^.

**Figure 3.**
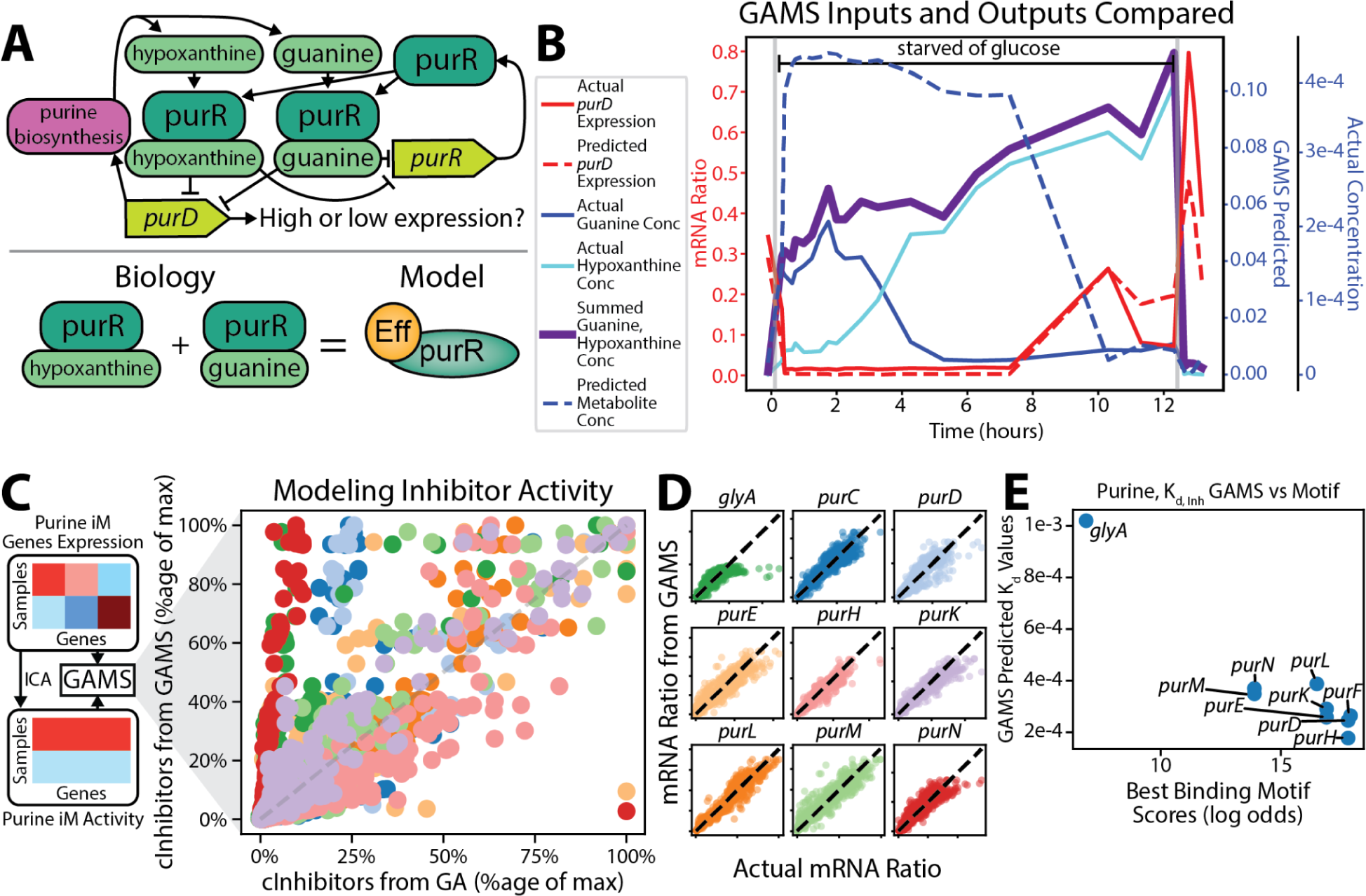
Model captures combined activity of multiple *PurR* effectors. **(A)** Regulatory network surrounding *purD* and *purR* involves two separate effector molecules which the products of *purD* and other genes produce. The underlying model treats both effectors as one pooled metabolite concentration. **(B)** Plotted actual and predicted values for metabolite concentrations and gene expression. The model initially accurately predicts both gene expression and metabolite concentration, but fails to accurately model metabolite concentration in the later stationary phase samples. **(C)** GAMS matches the measured and predicted inhibitor activity for the genes in the Purine iModulon. **(D)** The expression of the individual genes modeled in this iModulon can be accurately recreated. **(E)** The sequence-based predictions and model predictions for binding strength of *purR* to the genes it represses are negatively correlated.

Figure 3CD showcases how this model can accurately predict gene expression and recreates the input cInhibitor values for the majority of the genes. Similar to the Arginine case, the model’s predicted **K_d_** values also reliably anti-correlate with the sequence-based predictions of motif binding strength (Figure 3E).

### *Crp* ALE mutation prediction and validation

We next examined whether the modeling framework could accurately capture transcriptional regulation in cases of promoters affected by multiple transcription factors. We selected genes that appeared in both the Crp-2 iModulon and the Mlc iModulon. *ManXYZ* is activated by Crp-2 and repressed by DhaR, an iModulon with multiple primary regulators, one of which is *mlc*. *PtsG* is in the DhaR iModulon and is narrowly below the threshold for inclusion in the Crp-2 iModulon, so is included in Crp-2 for this study.

The model for this regulation and effectors involved is outlined in which both regulatory signals are required for accurate prediction of expression, as can be seen in Figure 4A. The expansion from one to two regulators requires the additional step of calculating sets of cActivator and cInhibitor that both satisfy the mathematical model and serve as good approximates of regulator activity. As described above, the extension of the workflow to multiple regulators is handled by a first step genetic algorithm which selects a set of valid solutions, which a greedy algorithm can further optimize in order to increase the correlation or anticorrelation between the cActivator or cInhibitor and the iModulon’s activity. Figure 4B and Figure 4C show these steps and how they generate input regulatory activities that satisfy the mathematical model that can be used in the GAMS solver. Figure 4DE shows how the GAMS model optimizes to both match the input cActivator and cInhibitor values as well as the expression values.

**Figure 4.**
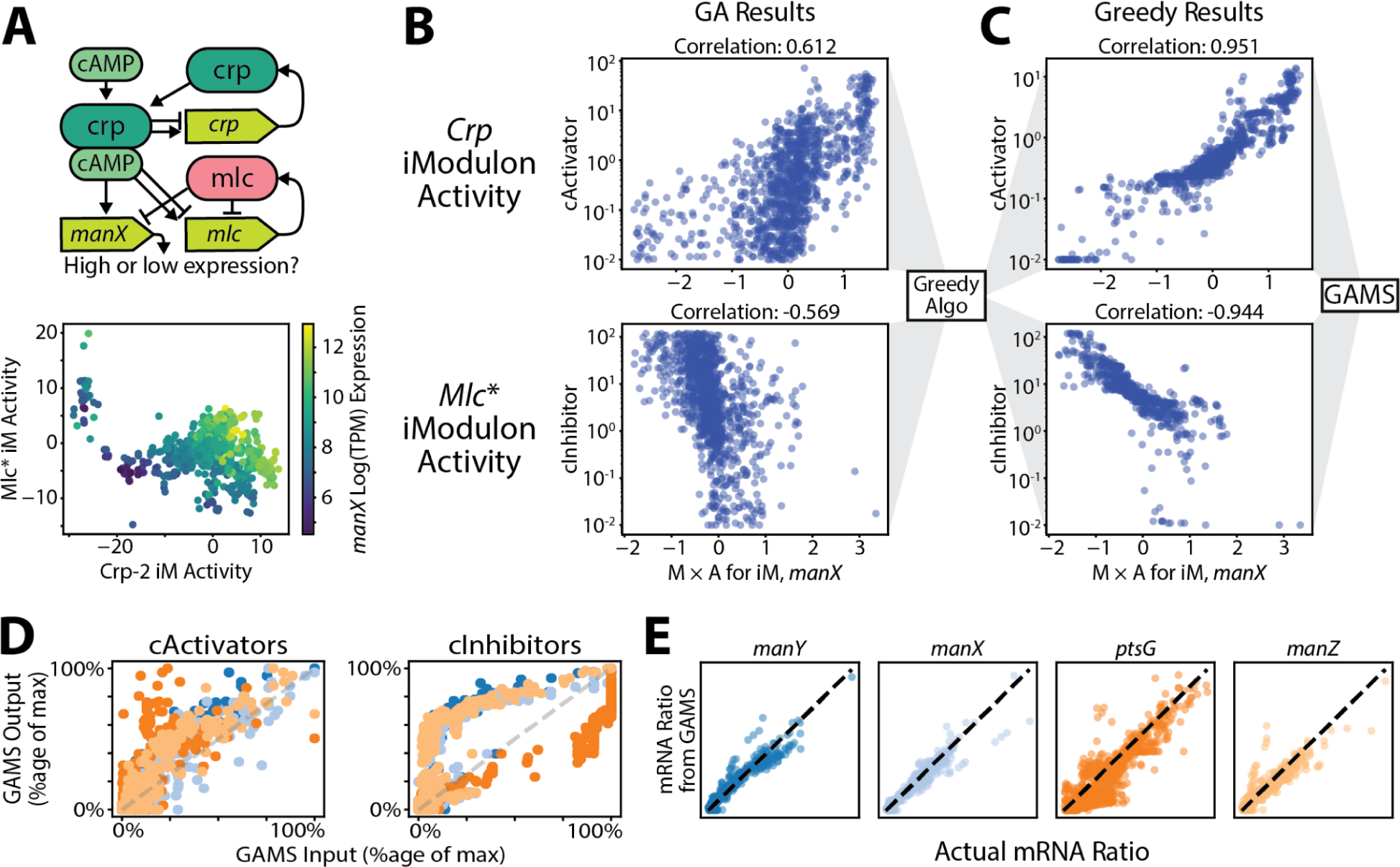
Models capture transcription factor interactions at Crp promoters. **(A)** The surrounding regulatory network of *manX* regulation which is repressed by *mlc* and promoted by *crp*, which also play roles in regulating themselves. The iModulons associated with these regulators determine *manX* expression. The Mlc iModulon is named DhaR in PRECISE1K as both *mlc* and *dhaR* are primary regulators of the same iModulon. **(B)** Because there is a promoter and inhibitor, there is one more unknown than equations and cActivator and cInhibitor must be calculated using a coupled genetic algorithm (GA). The resulting values are shown. **(C)** The results of the GA algorithm are then fed into a greedy algorithm which further optimizes the cActivator and cInhibitor values to be fed into GAMS. **(D)** The GAMS model matches the input cActivator and cInhibitor values from the greedy algorithm. **(E)** The GAMS model recreates the expression profiles of the modeled genes.

A *crp* knockout evolution experiment led to convergent mutations upstream of *ptsG* on repressor binding sites (NCBI GEO GSE266148). We tested our model’s ability to predict the impacts of these genetic alterations, by taking the control wildtype sample from said *crp* knockout evolution study and setting cActivator to zero to simulate the *crp* KO. Following this, cInhibitor was additionally set to zero to simulate the loss of the repressor. The predicted *crp* KO sample has lower expression than the predicted combined *crp* KO and repressor mutation sample. *Crp* KO strains from this study follow a similar pattern (see **Supplemental Figure 1**).

### Extending the workflow toward a quantitative reconstruction of the *E. coli* TRN

The high overlap between genes in iModulons and regulons enables the established workflow to use ICA-inferred component activities as a proxy for transcription factor regulatory activity (Figure 5A). Indeed, we observed that iModulons generally have higher correlations to gene expression than does expression of the regulator of the regulons to which a gene belongs (Figure 5B). Utilizing this high overlap, we have expanded our workflow to model twelve total regulatory cases. Five are inhibitor only (ArcA, Arginine, Cysteine-1, Fur-1, and Purine iModulons), six are promoter only (CpxR, Phosphate-1, Fur-2, Fnr-1, Cra, Crp-2, and SoxS), and one is dual promoter and inhibitor (Crp-2 and DhaR). In total, these cases account for 226 genes.

**Figure 5.**
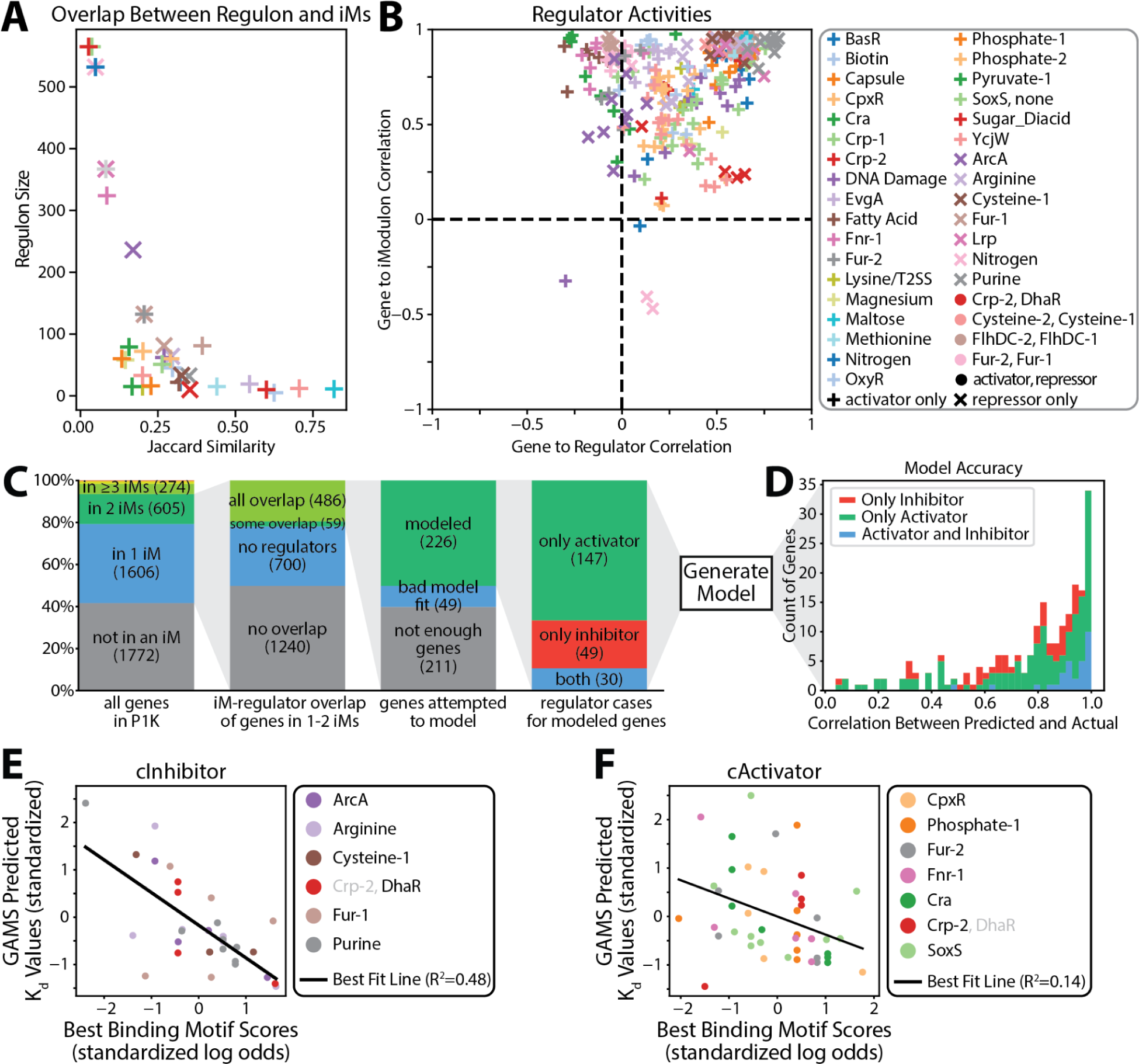
Scalability of the workflow toward capturing the E. coli transcription regulatory network. **(A)** The amount of overlap between iModulons and the regulons of the regulators of iModulons. High overlap exists for many iModulon-regulator pairs. **(B)** Correlation between genes and their iModulons and regulators. Genes have much higher correlation to their iModulons than regulators. **(C)** The various genes that are unable to be modeled are removed, the reasons for which are outlined. **(D)** The model’s accuracy across all samples is about equal for the different regulatory types. **(E)** The GAMS predicted **K_d_** values for the various inhibitors and the sequence-based predictions of motif binding strength are negatively correlated across all samples. Both values are standardized per regulator in order to scale them for comparison. **(F)** The GAMS predicted **K_d_** values for the various activators and the sequence-based predictions of motif binding strength are slightly negatively correlated across all samples. Both values are standardized per regulator in order to scale them for comparison.

Some genes are unable to be modeled for a variety of reasons, most often due to either not belonging to any iModulons or lack of annotated regulator of iModulons to which the gene belongs (Figure 5C). For the modeled genes, we are able to accurately recreate gene expression with a median correlation between predicted and actual of 0.82 (Figure 5D). The expected negative correlations between sequence-predicted motif scores and GAMS predicted **K_d_**values are found across the modeled regulons, although this component of the model’s accuracy is better for repressors than promoters (Figure 5EF). Thus, although the scope of the workflow remains somewhat limited compared to the entire *E. coli* TRN, the models that were able to be developed consistently perform well at capturing both gene expression and binding site strength.

## Discussion

Here, we developed a workflow to utilize ICA component (iModulon) activities extracted from gene expression databases as a proxy for regulator activities to parameterize transcriptional regulatory models in *E. coli*. Transcription regulation was modeled with standard thermodynamic binding equations based on well-established promoter complexes incorporating known transcription factor effectors. This workflow generates a data-backed physical model for gene regulation and enables the prediction of metabolites and transcription factor binding constants using expression data. These predictions can be verified using external data sources, thus showing the robustness of this workflow.

The Arginine iModulon, regulated by *argR*, serves as the primary repressor for several genes that control arginine import and biosynthesis. Expression of these genes is positively correlated with expression of their repressor *argR*, a counter-intuitive relationship that the model successfully captures by accounting for the role of the *argR* activator arginine, which governs the effective *argR* activity level. In another case study, *PurR*, which regulates the Purine iModulon, is activated by either guanine or hypoxanthine,^15^ and the model aggregates the effect of both effectors explicitly in its formulation. When compared to actual metabolite concentrations, the model is able to predict the sum concentration of both metabolites for the first few hours of a glucose starved dataset^14^. Thus, initial case studies suggest that the modeling formalism developed here can capture complex regulation scenarios. The lack of accuracy for the later hours is possibly due to our model not accounting for sigma factors which have a large effect on stationary phase expression^16^. Much effort has gone into modeling binding strength of sigma factors^17^ and proper utilization of this knowledge may enable its inclusion in a mechanistic model such as ours.

Many promoters are regulated by multiple transcription factors and other DNA-binding proteins. We developed a case study to examine the ability of the developed modeling formalism to capture regulator interactions at promoters. A small set of genes are promoted by the Crp-2 iModulon and inhibited by the DhaR iModulon, which are promoted by *crp* and *mlc* respectively. This regulatory case provides an example of our model predicting expression for genes with both an activator and repressor. The model is able to accurately model expression by providing well correlated cActivator and cInhibitor values through the use of genetic and greedy algorithms. Multi-regulator genes are common, as 1,413 genes have two or more annotated regulator binding sites^2^. To encompass these genes in future versions of this workflow, we will need to expand the number of different promoter states our model accounts for.

This workflow has currently been expanded to 12 regulatory cases which includes 226 genes. 54% of the genes in PRECISE1K account for only 20% of total expression variance^13^ and an additional 37% of the remaining genes have no annotated regulator^2^. This in total leaves 1,265 genes that explain significant variance and have known regulators, of which 138 are included in our model. To further expand this model, we will need to include genes not currently in iModulons or include the genes of iModulons that are not directly regulated by known iModulon regulators. Larger-scale models can account for more genes^18^, but lack the mechanical details that a workflow such as ours allows. Combining these approaches could potentially yield a detailed mechanistic model for all of transcription.

The mechanistic promoter models presented here connect together detailed biological research about promoter-regulator interaction with the global regulatory trends revealed through large-scale data analytics. This enables the inferrance of biological constants and metabolite measurements which can be directly mapped to gene expression. Whole-cell models exist which are able to connect metabolism to the cell’s environment through the utilization of differential equations and numerous parameters^8,19^. The promoter regulation models developed here could presumably be integrated within multi-scale and whole cell frameworks to better represent transcriptional regulation during life cycle simulations. Taken together, this study suggests a path toward generating a large-scale mechanistic model of gene regulation in *E. coli*.

## Methods

### iModulon Computation

RNA-sequencing data was used to create iModulon activity levels of our strains using PyModulon^12^ which is available at https://github.com/SBRG/pymodulon. Activities of iModulons were compared to samples from PRECISE1K which is easily accessible using iModulonDB^20^. The calculations of new iModulons for our dataset in addition to PRECISE1K were performed using iModulonMiner^21^ which is available at https://github.com/SBRG/iModulonMiner. iModulonDB (https://imodulondb.org/^20^) for a more complete description of iModulons and their calculation.

### Workflow Overview

The process described here is outlined in **Supplemental Figure 2**. Before any data is processed, genes are selected for modeling (see **Methods: Initial Selection of Genes to Model**) and outliers samples are removed (see **Methods: Removal of Outlier Samples**). The initial inputs are gene expression files in log TPM format, the M and A matrices output from ICA, and a proteomic dataset^22^. These three datasets are used to create mRNA expression ratios, recreated gene expression values from M × A, and creating constraints for the model parameters. Independent of this, ideal biological constants, namely **K_d, RNAP_**, **k_escape_**, and **K_eq, opening_** are selected from a calculated set of possible solutions to the mathematical equations (see **Methods: Selection of Biological Constants**). The M × A values for each gene and these ideal biological constants are used to generate cActivator and cInhibitor values (see **Methods: Creating cActivator and cInhibitor Values**). If there is both an activating and inhibiting iModulon, a genetic algorithm and following greedy algorithm are used to further improve the cActivator and cInhibitor values (see **Methods: GA and Greedy Algorithm Optimization**).

These cActivator and/or cInhibitor values are then input into the GAMS optimization software which recreates new cActivator and cInhibitor values calculated from the correct regulator type equation (see **Methods: Physical Model Equations**). GAMS optimizes two primary criteria, matching to the input cActivator and/or cInhibitor values and matching gene expression values to the actual gene expression values. The GAMS model then outputs the underlying constants for the new cActivator and cInhibitor values as well as the predicted gene expression values (see **Methods: GAMS Model**). The GAMS model can be rerun multiple times with changing constraints and weightings in order to achieve higher agreement between output and input values (see **Methods: GAMS Parameter Optimization**).

### Initial Selection of Genes to Model

In order for a gene to be included in our transcriptional model, it must satisfy several criteria. First, the gene must be in one or two iModulons that have a well characterized regulator that also regulates the gene. This regulation is determined by the existence of a strong or confirmed regulatory relationship according to RegulonDB^2^. If a gene is in two iModulons, it must be repressed by one and promoted by the other.

At this point, initial biological constants can be generated for the gene and produce cActivator and cInhibitor values. For some genes, there is no valid set of biological constants which result in usable cActivator or cInhibitor values. This is most often due to either extremely high or low expression values for the gene without either violating the underlying mathematical equations or creating negative cActivator or cInhibitor values. The transcription factors themselves, often members of their own iModulons, are also removed. All remaining genes are grouped by their respective iModulons, into their regulatory cases: activator only cases, inhibitor only cases, or dual activator and inhibitor cases After these steps, regulatory cases are removed if they contain only one sample.

### Mathematical Model Equations

The basis for the transcriptional regulatory model is a mass balance reaction describing mRNA transcription and degradation/dilution, and standard thermodynamic binding equations^23^ describing RNA polymerase binding, formation of a transcription bubble and escape from the promoter complex, as well as transcription factor binding that either activates or inhibits RNAP binding. Resulting equations are outlined below:

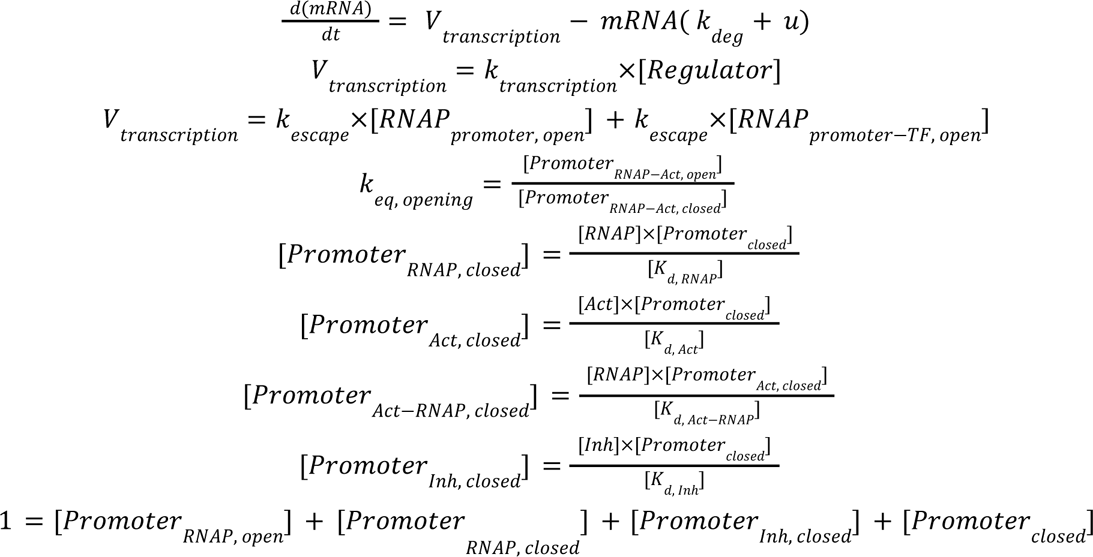

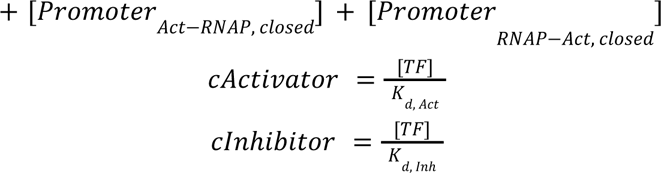

The above equations are used for calculating mRNA as a function of regulatory activity according to the following formula:

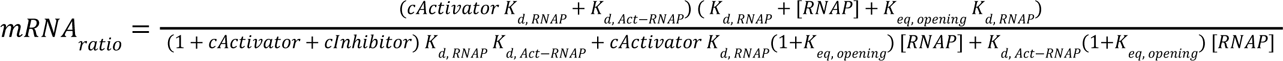

To incorporate different effector binding types, the formula for cActivator and cInhibitor have a few various forms. The following equations are used for calculating cActivator or cInhibitor in terms of metabolite concentrations and **K_d, Act_** in the case that the transcription factor has multiple (3) effector binding sites:

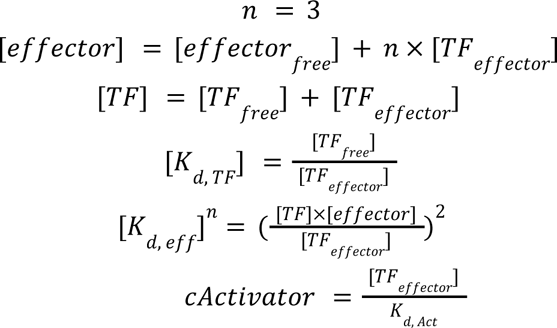

Solving for cActivator yields with n = 3 (cInhibitor has the same form):

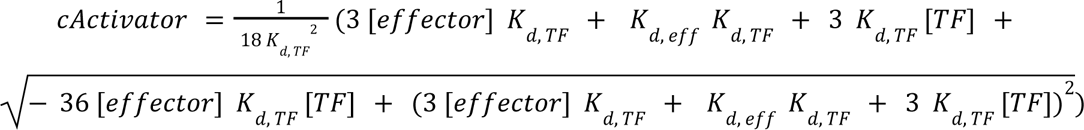

The following equations are used for calculating cActivator in terms of metabolite concentrations and **K_d, Act_** in the case that the transcription factor has n co-effectors:

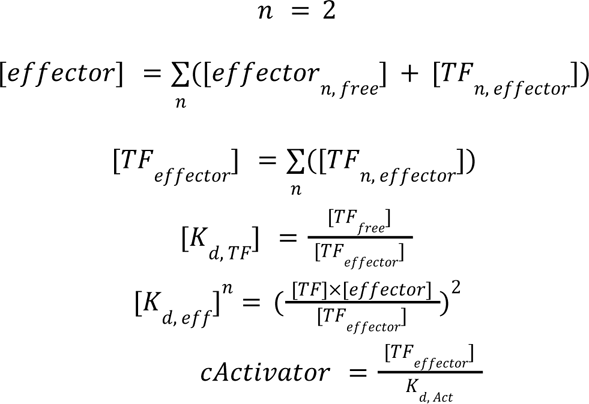

Solving for cActivator yields with n = 2 (cInhibitor has the same form):

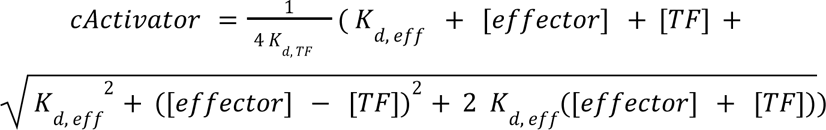

For the regulatory types involving effectors, TF concentration is set for each individual sample by scaling the overall expression values of the TF to be between the minimum and maximum of a proteomics dataset^22^.

### Removal of Outlier Samples

As the GAMS model uses squared difference regression objectives, outliers can have large effects on the outcome by setting either extremely high or extremely low cActivator or cInhibitor values. To avoid this, a few outlier samples are typically removed for every regulatory case. For each regulatory case, a correlation matrix is created between the samples across the regulatory case’s genes. Any sample that does not have greater than or equal to 0.5 correlation value to at least 5% of the other samples is removed. For all regulatory cases, this is always under 4% of total samples.

### Selecting Basal Conditions and Conversion to mRNA Ratios

In order to eliminate variables from the equations, mRNA ratios are used as opposed to log TPM expression values. In order to calculate these values, a basal condition must be selected. This condition must be uniform across each regulatory case. For each regulatory case, the expression profiles for the genes of said case are collected and standardized, the resulting distributions are then used to select the basal condition. If the regulatory case is only an inhibitor, the sample with the highest average standardized expression is chosen. If the regulatory case is only an activator, the sample with the lowest average standardized expression is chosen. In a dual promoter and inhibitor case, the sample with the closest to zero average standardized expression is chosen.

All expression values are then un-logged and divided by the un-logged value for their respective basal condition. This generates an mRNA ratio value. The same un-logging and dividing by the basal condition is performed on the ICA reconstructed expression values (M × A) in order to scale them properly so that they can be used to calculate the cInhibitor or cActivator values in the non dual-regulator cases.

### Selection of Biological Constants

Solving a steady state transcription rate and substituting this expression into the mass balance equation for mRNA, as described above, result in the following equations for steady-state mRNA concentration:

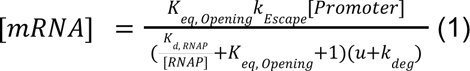

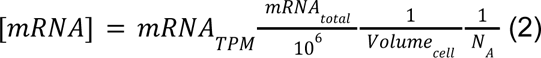

The following constants in the above equations are defined as follows:

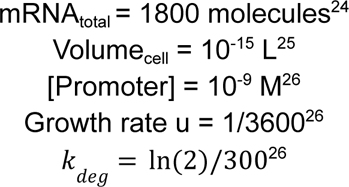

A grid of possible solutions for **K_d, RNAP_**, **k_escape_**, and **K_eq, opening_**is created which is later tested to determine a best set of values for the model for each gene. First a range of possible values are assumed for **K_d, RNAP_** = 10^[-7,-5]^ and **k_escape_** = 10^[-3, 1]^. We calculate the minimum viable value for **k_escape_** assuming the max **K_eq, opening_** = 100 and update the range of possible values for **K_d, RNAP_** and **k_escape_**. We generate nine new pairwise combinations of **K_d, RNAP_** and **k_escape_**which satisfies the equation. The value of **K_eq, opening_** is calculated in order to solve the equation when [mRNA] is known. This process altogether creates 9 valid sets of constants which correctly calculate [mRNA].

**K_d, Act-RNAP_** is set through a two step optimization process for any regulator case with an activating iModulon. In the first step, **K_d, Act-RNAP_** is set to **K_d, RNAP_** which creates negative cActivator values. **K_d, Act-RNAP_** is then incrementally reduced until there are no longer any negative cActivator values. This sets the maximum value for **K_d, Act-RNAP_**. The second step improves the distribution of cActivator values to both be between 0 and 1,000 and more evenly spread by maximizing the 80th percentile value of the cActivator values. This is to prevent the creation of extreme outlier cActivator values which GAMS will then overfit on if not corrected.

The selection of **K_d, Act-RNAP_** described above is carried out for each of these 9 sets of valid constants and the set with the lowest **K_d, RNAP_** which also is able to create a valid **K_d, Act-RNAP_** is selected for the gene. For some genes, there is not a valid **K_d, Act-RNAP_** value which satisfies the equation without creating negative cActivator values, in which case the gene is removed from the model.

### Creating cActivator and cInhibitor Values

If there is only a single activator (cActivator) or inhibitor (cInhibitor) for a given promoter, the same number of unknown variables and equations exist so the value can be set. Instead of directly inputting the mRNA ratio value, we input ICA reconstructed expression values (M × A) in order to better model regulatory activity as opposed to expression. These cActivator or cInhibitor values are then calculated for each sample in the gene and each gene in the regulatory case.

If there is both an activator and inhibitor at a given promoter, the cActivator and cInhibitor values need to be calculated simultaneously. First a set of valid solutions is found with various values of cInhibitor and cActivator, initially ranging between 0 and 1,000. These valid solutions are then passed to the genetic algorithm to select sets that correlate with the inhibiting and activating iModulons.

### Genetic and Greedy Algorithm Optimization

If there is both an activator and inhibitor at a given promoter (resulting in both (cActivator and cInhibitor needing to be parameterized), the underlying mathematical equations must be solved simultaneously for both cActivator and cInhibitor. To pick valid values of cActivator and cInhibitor that also correlate with their respective iModulon activities and thus regulatory activity, two algorithms are used.

First, a Genetic Algorithm (GA) gives us an initial solution^27^ using a modified version of eaMuPlusLambda. The GA used for this study is multi-objective as it attempts to both maximize the Spearman correlation between the cActivator and the activating iModulon’s activity and minimize the Spearman correlation between cInhibitor and the inhibiting iModulon’s activity (as we want a highly negative correlation for cInhibitor and the inhibitor iModulon). Each individual in the GA consists of a randomly-sampled valid solution for cActivator and cInhibitor for every condition. A population of these individuals is created by random individual creation 100 times. Each individual’s fitness is evaluated using the two objectives and ranked based on their performance. The best individuals from each generation are selected to propagate to the next generation using the SPEA2 algorithm^28^. 5% of the selected individuals undergo mutation in which a random cActivator-cInhibitor pair is selected for a random amount of conditions in the individual. An additional 5% of the selected individuals undergo crossover in which a random condition’s cActivator-cInhibitor pair is swapped between two individuals.

The values generated by this algorithm do drastically improve the positive and negative correlations for cActivator and cInhibitor, but an additional greedy algorithm further optimizes the solution. This algorithm searches the local cActivator-cInhibitor pairs to find better ones using local gradient information. The order of the conditions are randomly shuffled for the best individual in the current generation. Each other condition is iterated over to see if there is a better scored cActivator-cInhibitor pair within 10 steps. This process is then carried out for each condition. The entire greedy algorithm is repeated 50 times. This results in highly correlated and negatively correlated cActivator and cInhibitor values that are also valid solutions to the underlying mathematical equations.

### Selection of Constraints on the GAMS Model

The variables that GAMS uses to calculate cActivator and cInhibitor (and thus gene expression) are initially constrained by experimentally derived data. For K_d, Act_ and K_d, Inh_ this constraint is initially based experimentally derived constants for *crp^29^*. The initial constraints are the minimum observed K_d_ value for *crp* divided by 1,000 and the maximum observed K_d_ value for *crp* multiplied by 1,000 times. The metabolite concentrations are constrained by the minimum observed concentration divided by 1,000 and the maximum observed concentration multiplied by 1,000^14^. If no such measurement of the metabolite exists in *Link et al. 2015*, the minimum of any metabolite in said study divided by 1,000 is used as the minimum and the maximum of any metabolite in said study multiplied by 1,000 is used as the maximum.

### GAMS Model

The GAMS workflow is a multi-parameter optimization model that optimizes two objectives: 1) matching actual and predicted mRNA ratios, and 2) matching actual and predicted regulator activities. The first step is reading in of constants and constraints for modeled parameters, which are previously generated as well as a weighting factor to balance between the two objectives. The input cActivator and cInhibitor values are also input, along with a mapping of what type of regulator each iModulon is modeled as. The model variables are created, constrained, and populated with initial values.

The dnlp solver is used for our model. The optimization is then run to minimize the squared error of both objective functions. Once complete, the following modeled variables are output: predicted mRNA ratio, cActivator, cInhibitor, and the various variables that cActivator and cInhibitor are calculated based on. Depending on the regulator type, these variables are some combination of effector concentrations, regulator concentrations, **K_d, Act_**, and **K_d, Inh_**.

### GAMS Parameter Optimization

A few important constrained variables and the weighting between the two criteria have a large impact on model performance. These constrained variables, depending on the regulator type, are effector concentrations, regulator concentrations, **K_d, Act_**, and **K_d, Inh_**. In order to pick constraints that lead to more accurate models while still maintaining biologically-relevant constraints, we have developed a parameter optimization pipeline.

This pipeline creates a set of values for each of these important inputs to GAMS. This set contains one value that is equal to the default value, another that is X% higher, and a third that is X% lower with X starting at 200. GAMS is then run on each combination of these sets (so N^3^ runs where N is how many parameters are tested). The resulting run with the least squared error for the objective functions is chosen. If there is a tie, usually due to loose constraints, the run with the tightest constraints is chosen. The process above is repeated with the values from the chosen run now being the default values and X being scaled down by 20% of its previous value. This process is repeated until no more large reductions in the error are found.

### Motif Analysis

The motif analysis was performed using https://github.com/SBRG/bitome^4^. The motif_search function was used for each using regulonDB as a source for finding the ideal binding motif to scan with. For each transcription factor, each of its regulated genes were scanned from 100 bp upstream to 50 bp downstream for a matching binding motif.

## Acknowledgements

This work was supported by the Novo Nordisk Foundation [grant number NNF20CC0035580]. We would like to thank Archana Balasubramanian for helpful discussions on this manuscript.

## Supplement

**Supplemental Figure 1.**
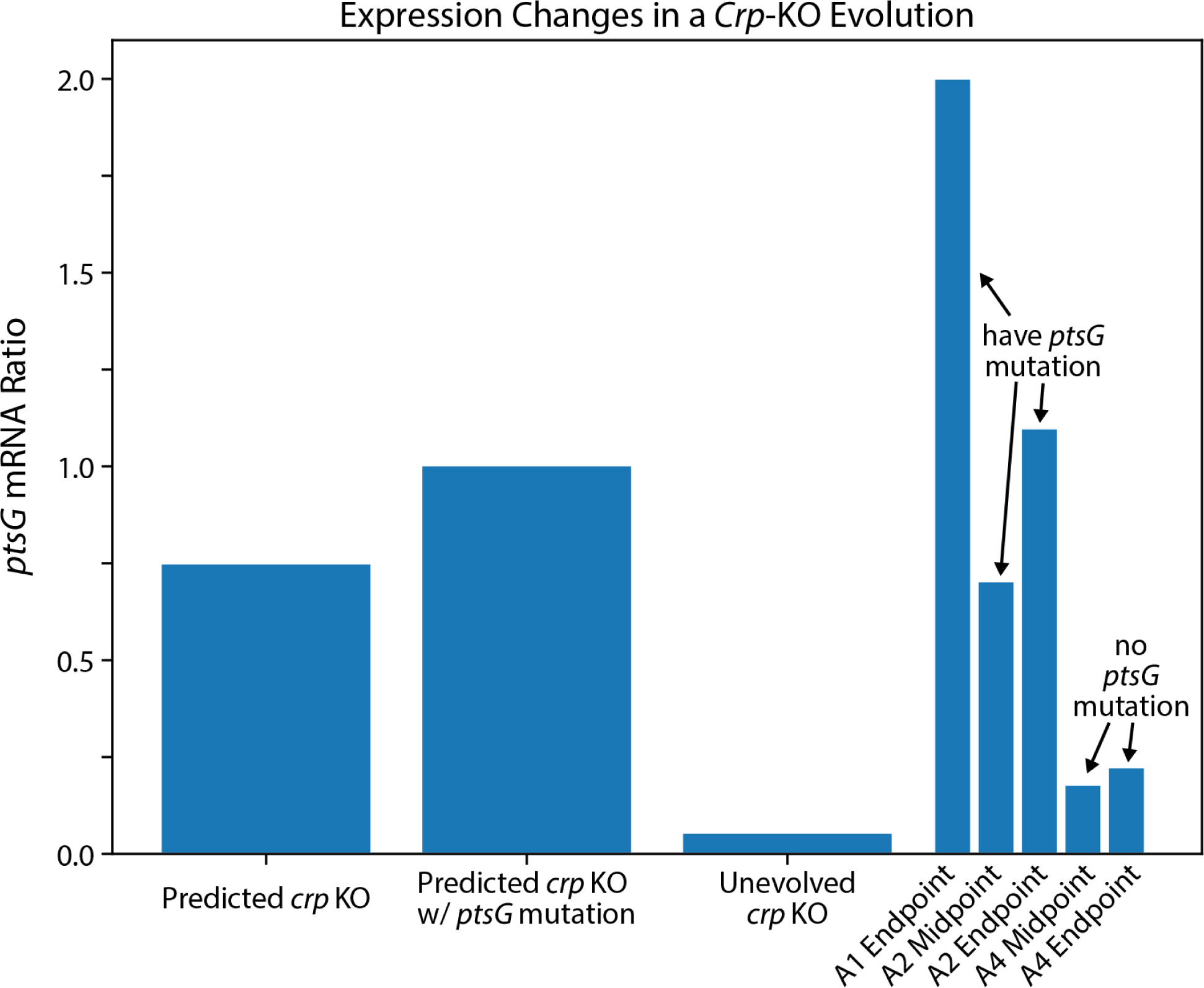
Modeling of expression changes caused by regulatory mutations. A *crp* KO evolution study led to convergent mutations of repressor sites upstream of *ptsG* (NCBI GEO GSE266148). Our model predicts higher expression with an upstream *ptsG* mutation. A similar trend can be seen in the experiment’s data.

**Supplemental Figure 2.**
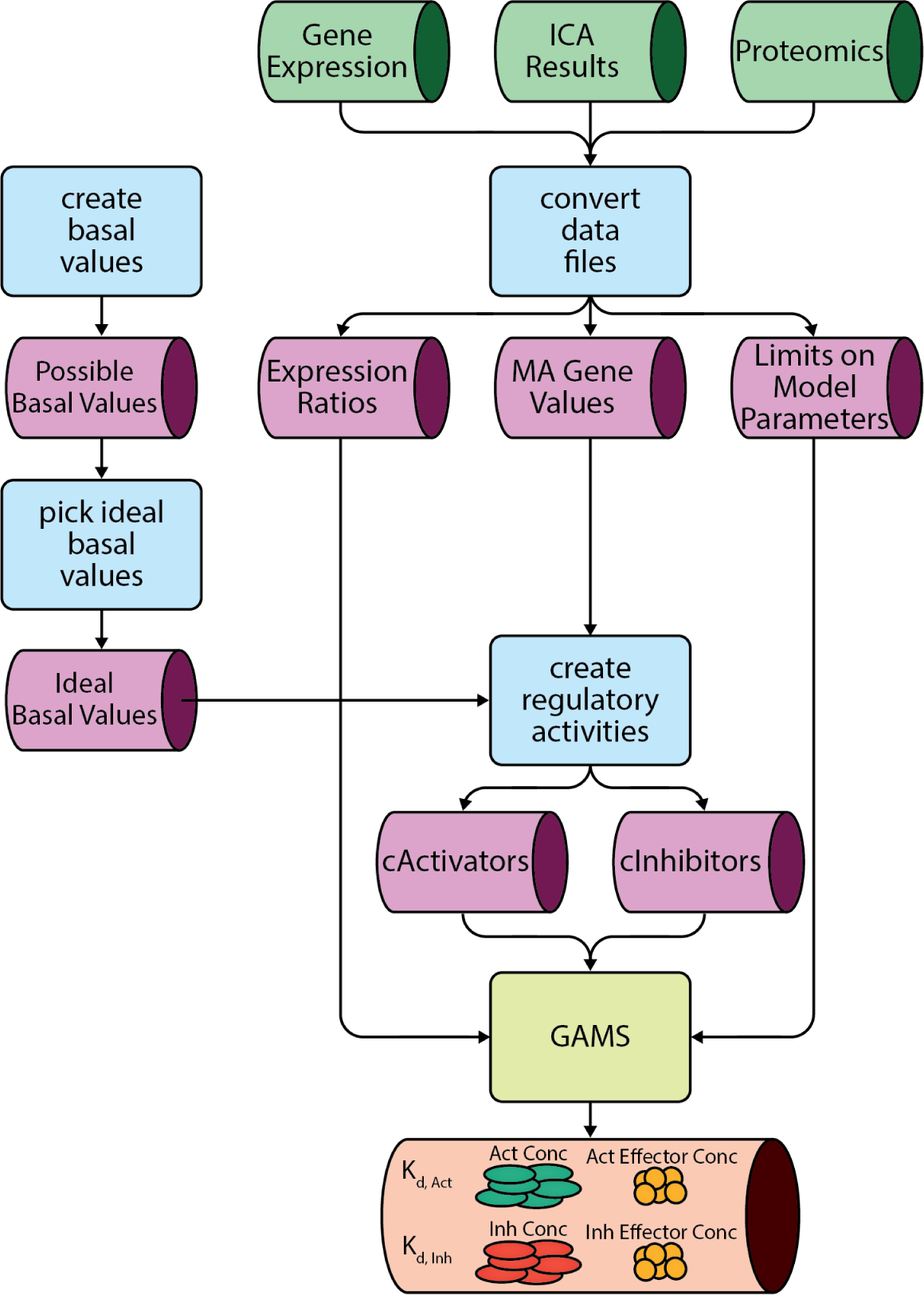
Outline of the model’s workflow. The tubes represent data while the squares represent code. Green represents input data, purple are internal data types used within the workflow, and red represents output data. Blue code boxes are run for every gene while GAMS is run for all genes of a regulatory case.

## References

1. Bervoets, I. & Charlier, D. Diversity, versatility and complexity of bacterial gene regulation mechanisms: opportunities and drawbacks for applications in synthetic biology. FEMS Microbiol. Rev. 43, 304–339 (2019).

2. Santos-Zavaleta, A. et al. RegulonDB v 10.5: tackling challenges to unify classic and high throughput knowledge of gene regulation in E. coli K-12. Nucleic Acids Res. 47, D212–D220 (2018).

3. Madan Babu, M. & Teichmann, S. A. Evolution of transcription factors and the gene regulatory network in Escherichia coli. Nucleic Acids Res. 31, 1234–1244 (2003).

4. Lamoureux, C. R. et al. The Bitome: digitized genomic features reveal fundamental genome organization. Nucleic Acids Res. 48, 10157–10163 (2020).

5. Edgar, R., Domrachev, M. & Lash, A. E. Gene Expression Omnibus: NCBI gene expression and hybridization array data repository. Nucleic Acids Res. 30, 207–210 (2002).

6. Marbach, D. et al. Wisdom of crowds for robust gene network inference. Nat. Methods 9, 796–804 (2012).

7. Feist, A. M., Herrgård, M. J., Thiele, I., Reed, J. L. & Palsson, B. Ø. Reconstruction of biochemical networks in microorganisms. Nat. Rev. Microbiol. 7, 129–143 (2009).

8. Covert, M. W., Knight, E. M., Reed, J. L., Herrgard, M. J. & Palsson, B. O. Integrating high-throughput and computational data elucidates bacterial networks. Nature 429, 92–96 (2004).

9. Cho, B.-K. et al. The transcription unit architecture of the Escherichia coli genome. Nat. Biotechnol. 27, 1043–1049 (2009).

10. Bailey, T. L. et al. MEME SUITE: tools for motif discovery and searching. Nucleic Acids Res. 37, W202–8 (2009).

11. Karp, P. D. et al. The EcoCyc Database. EcoSal Plus 8, (2018).

12. Sastry, A. V. et al. The Escherichia coli transcriptome mostly consists of independently regulated modules. Nat. Commun. 10, 5536 (2019).

13. Lamoureux, C. R. et al. A multi-scale expression and regulation knowledge base for Escherichia coli. Nucleic Acids Res. 51, 10176–10193 (2023).

14. Link, H., Fuhrer, T., Gerosa, L., Zamboni, N. & Sauer, U. Real-time metabolome profiling of the metabolic switch between starvation and growth. Nat. Methods 12, 1091–1097 (2015).

15. Choi, K. Y. & Zalkin, H. Structural characterization and corepressor binding of the Escherichia coli purine repressor. J. Bacteriol. 174, 6207–6214 (1992).

16. Hirsch, M. & Elliott, T. Role of ppGpp in rpoS stationary-phase regulation in Escherichia coli. J. Bacteriol. 184, 5077–5087 (2002).

17. Rhodius, V. A. & Mutalik, V. K. Predicting strength and function for promoters of the Escherichia coli alternative sigma factor, σE. Proceedings of the National Academy of Sciences 107, 2854–2859 (2010).

18. Macklin, D. N. et al. Simultaneous cross-evaluation of heterogeneous datasets via mechanistic simulation. Science 369, (2020).

19. Kotte, O., Zaugg, J. B. & Heinemann, M. Bacterial adaptation through distributed sensing of metabolic fluxes. Mol. Syst. Biol. 6, 355 (2010).

20. Rychel, K. et al. iModulonDB: a knowledgebase of microbial transcriptional regulation derived from machine learning. Nucleic Acids Res. 49, D112–D120 (2021).

21. Sastry, A. V. et al. Mining all publicly available expression data to compute dynamic microbial transcriptional regulatory networks. bioRxiv 2021.07.01.450581 (2021) doi:10.1101/2021.07.01.450581.

22. Schmidt, A. et al. The quantitative and condition-dependent Escherichia coli proteome. Nat. Biotechnol. 34, 104–110 (2016).

23. Einav, T. & Phillips, R. How the avidity of polymerase binding to the –35/–10 promoter sites affects gene expression. Proceedings of the National Academy of Sciences 116, 13340–13345 (2019).

24. Moran, M. A. et al. Sizing up metatranscriptomics. ISME J. 7, 237–243 (2012).

25. Kubitschek, H. E. & Friske, J. A. Determination of bacterial cell volume with the Coulter Counter. J. Bacteriol. 168, 1466–1467 (1986).

26. Milo, R., Jorgensen, P., Moran, U., Weber, G. & Springer, M. BioNumbers--the database of key numbers in molecular and cell biology. Nucleic Acids Res. 38, D750–3 (2010).

27. De Rainville Université Laval Québec PQ Canada, F.-M., Félix-Antoine Fortin Université Laval, Québec, PQ, Canada, Marc-André Gardner Université Laval, Québec, PQ, Canada, Marc Parizeau Université Laval, Québec, PQ, Canada & Christian Gagné Université Laval, Québec, PQ, Canada. DEAP. https://dl.acm.org/doi/10.1145/2330784.2330799 doi:10.1145/2330784.2330799.

28. Zitzler, E., Laumanns, M. & Thiele, L. SPEA2: Improving the strength pareto evolutionary algorithm. TIK Report 103, (2001).

29. Gunasekara, S. M. et al. Directed evolution of the Escherichia coli cAMP receptor protein at the cAMP pocket. J. Biol. Chem. 290, 26587–26596 (2015).

30. Browning, D. F. & Busby, S. J. W. The regulation of bacterial transcription initiation. Nat. Rev. Microbiol. 2, 57–65 (2004).

31. Martínez-Antonio, A., Janga, S. C. & Thieffry, D. Functional organisation of Escherichia coli transcriptional regulatory network. J. Mol. Biol. 381, 238–247 (2008).

32. Meysman, P. et al. Use of structural DNA properties for the prediction of transcription-factor binding sites in Escherichia coli. Nucleic Acids Res. 39, e6 (2011).

33. Ireland, W. T. et al. Deciphering the regulatory genome of Escherichia coli, one hundred promoters at a time. Elife 9, e55308 (2020).

34. Myers, K. S., Park, D. M., Beauchene, N. A. & Kiley, P. J. Defining bacterial regulons using ChIP-seq. Methods 86, 80–88 (2015).

35. Federowicz, S. et al. Determining the Control Circuitry of Redox Metabolism at the Genome-Scale. PLoS Genetics vol. 10 e1004264 Preprint at 10.1371/journal.pgen.1004264 (2014).

36. Latif, H. et al. ChIP-exo interrogation of Crp, DNA, and RNAP holoenzyme interactions. PLoS One 13, e0197272 (2018).

37. Cho, B.-K., Barrett, C. L., Knight, E. M., Park, Y. S. & Palsson, B. Ø. Genome-scale reconstruction of the Lrp regulatory network in Escherichia coli. Proc. Natl. Acad. Sci. U. S. A. 105, 19462–19467 (2008).

38. De Smet, R. & Marchal, K. Advantages and limitations of current network inference methods. Nat. Rev. Microbiol. 8, 717–729 (2010).

39. Hyvärinen, A. & Oja, E. Independent component analysis: algorithms and applications. Neural Networks vol. 13 411–430 Preprint at 10.1016/s0893-6080(00)00026-5 (2000).

40. Rychel, K., Sastry, A. V. & Palsson, B. O. Machine learning uncovers independently regulated modules in the Bacillus subtilis transcriptome. Nat. Commun. 11, 6338 (2020).

